# Viral and bacterial plant pathogens suppress antiviral defense against flaviviruses in their insect vectors

**DOI:** 10.1101/2024.08.15.608128

**Authors:** Adriana Larrea-Sarmiento, Alejandro Olmedo-Velarde, Michael West-Ortiz, Douglas Stuehler, Saeed Hosseinzadeh, Aminah Coleman, Stephanie Preising, Glenn Parker, Zhangjun Fei, Michelle Heck

**Author notes:** Corresponding author: Michelle Heck. Department of Plant Pathology, Entomology and Microbiology, Iowa State University, Ames, IA USA.

## Abstract

A positive, single-stranded RNA virus member within the *Flavivirus* genus was identified and characterized infecting *Myzus persicae*. This new insect-specific virus (ISV), Myzus persicae flavivirus (MpFV), is 23,236 nucleotides in length and encodes a large polyprotein from a single open reading frame. Analysis of conserved domains showed that helicases, NS3-proteases, Fts-J methyltransferase, and an RNA-dependent RNA polymerase are present in the coded polyprotein. Aphid-infecting ISVs have been reported to interact with plant viruses within the vector, modulating its titer and manipulating aphid behavior and morphology. Small RNA (sRNA) profile analysis of the *M. persicae* sRNA profile demonstrated that the circulative plant virus, potato leafroll virus (PLRV), modified the aphid antiviral immunity against MpFV. Abundant sRNA reads matching MpFV were detected when aphids were fed on healthy plants, sucrose diet, and potato virus Y-infected plants. In contrast, no MpFV reads were detected in aphids that had acquired PLRV from infected plants or artificial diet sachets containing purified virions. While the titer of *M. persicae densovirus* (MpDNV) was previously reported to be regulated by expression of the PLRV silencing suppressor protein P0, P0 had no effect on MpFV titer in the aphid. MpFV was transmitted 100% vertically to the offspring, and exhibited tissue tropisms for the body rather than the head. By artificial diet assays, other aphid species, including *Aphis gossypii* (cotton aphid), *Schizaphis graminum* (greenbug aphid), *Rhopalosiphum* padi (bird cherry-oat aphid), and *R. maidis* (corn leaf aphid), acquired the MpFV. These findings further support the idea that PLRV suppresses aphid immunity against ISVs, suggest the existence of at least two distinct pathways for PLRV-induced aphid immune system modulation. To test whether other circulative plant pathogens suppress insect anti-viral immunity against insect-specific flaviviruses, we quantified the small RNA response of *Diaphorina citri*, vector of “*Candidatus* Liberibacter asiaticus” (*C*Las) associated with citrus greening disease and showed that *C*Las also suppresses *D. citri* anti-viral immunity against *D. citri-like flavivirus* (DcLFV). These data reveal an evolutionary conserved, unexpected role for diverse circulative plant pathogens in modulating anti-viral immunity in hemipteran vectors.

## INTRODUCTION

The green peach aphid, *Myzus persicae*, is a polyphagous, worldwide hemipteran pest capable of transmitting over 100 plant viruses. Transmission of plant viruses by aphids involves co-evolved, multitrophic interactions among plant viruses, aphid vectors, and plant hosts (Castle et al. 1998; Gray et al. 2014; Pinheiro et al. 2019; Ray and Casteel 2022). In potato (*Solanum tuberosum*), the green peach aphid can cause yield losses by directly feeding on phloem sap and, more significantly, by transmitting plant viruses that include the circulative, non-propagative potato leafroll virus (PLRV; Genus *Polerovirus*, Family *Solemoviridae*). In the laboratory, several solanaceous plant species are used as hosts to rear and study *M. persicae*. *Nicotiana benthamiana*, *Physalis floridana* (downy ground cherry), and *Brassica napa* (turnip) serve as model plants for studying the transmission of the circulative PLRV and the non-persistently transmitted potato virus Y (PVY), using different lineages of *M. persicae* (Lee et al. 2002; Pinheiro et al. 2019; Ramsey et al. 2007).

In addition to transmission of plant viruses, aphids are known to be a host of several insect-specific viruses (ISVs). The wide use of high-throughput sequencing (HTS) has led to the discovery of DNA and RNA ISVs infecting aphids within the families *Flaviviridae* and *Parvoviridae* and nege-, kita-, picorna-like viruses have been reported (Kondo et al. 2020; Pinheiro et al. 2019; Teixeira et al. 2016; van der Wilk et al. 1997; van Munster et al. 2003) (Moon et al. 1998; Ryabov 2007; Ryabov et al. 2009; van Munster et al. 2002).

The family *Flaviviridae* contains virus members that can be divided into two broad groups based on their host range: (i) viruses that infect vertebrates and their insect vector and (ii) those that infect only invertebrates. Dual-host flaviviruses that infect insects and vertebrates typically undergo replicative cycles in both their insect and vertebrate host and include important human and veterinary pathogens, including dengue viruses, Zika virus, and bovine viral diarrhea viruses. The second group of RNA insect viruses are the ISVs, infecting insects and insect cells, but they do not replicate in vertebrates or vertebrate cell lines (Kuno 2004, 2007; Vasilakis and Tesh 2015). A third group includes vertebrate-only flaviviruses, also known as the ‘no known vector’ group, that naturally infect and are transmitted between vertebrates but neither have known arthropod association nor infect insect cell lines (Kuno 2004).

Flaviviruses have a positive-sense, single-stranded RNA genome of ∼ 10-11 kb encoding one single open reading frame (ORF) that is flanked by the 5 and 3 untranslated regions (UTR). The long polyprotein is cleaved by viral and host proteases during replication into 10 mature proteins, three structural proteins: capsid (C), pre-membrane/membrane (prM/M), and envelope (E), and seven non-structural (NS) proteins: NS1, NS2A, NS2B, NS3, NS4A, NS4B, and NS5. The 5 UTR is capped and the 3 UTR is highly structured non-coding that coordinates viral translation, replication and host immune modulation. Moreover, the 3 UTR is involved in the generation of subgenomic flavivirus RNA, helping the virus to evade the host immune response (Ng et al. 2017; Paraskevopoulou et al. 2021). Although ‘large genome flaviviruses’ (LGFs) with genome lengths ranging from ∼ 16-23 kb have been reported lately infecting psyllids, bees, flies, nematodes, aphids, as well as plants, their genome presents a typical flavivirus genome organization (Rashidi et al. 2022; Kondo et al. 2020; Paraskevopoulou et al. 2021; Teixeira et al. 2016).

Myzus persicae densovirus (MpDNV; genus *Ambidensovirus*, family *Parvoviridae*), a single-stranded DNA virus, has been characterized to interact with PLRV within the aphid during co-infection. On potato, as the virus infection progresses, the PLRV viral suppressor protein, P0 (P0), inhibits the plant’s silencing defense mechanism via degradation of the plant ARGONAUTE1 (AGO1) mediated by the F-box-like domain through an autophagy pathway (Barrios Barón et al. 2021; Fusaro et al. 2012; Michaeli et al. 2019). Published work from our group has shown that once the plant immune system is turned down by P0, the aphids anti-viral RNA interference immune response against MpDNV is also turned down, resulting in PLRV-viruliferous aphids with a higher titer of MpDNV (Pinheiro et al. 2019). Transient expression of P0 in the plant raises MpDNV titer, which has been associated with increased alate production and aphid fecundity, suggesting that both plant and insect viruses manipulate aphid biology to favor viral spread (Pinheiro et al. 2019). Although this is an indirect effect of MpDNV on PLRV transmission, little is known about the effects of ISVs during plant virus acquisition and transmission.

In the present study, we report the molecular and biological characterization of a new ISV infecting the green peach aphid. The name Myzus persicae flavivirus (MpFV) is proposed for this novel virus that should be a member within an unclassified cluster of ISV within the family *Flaviviridae,* that shares similarities with other LGFs. Experiments demonstrate that PLRV infection in plants modulates MpFV titer in the aphid. Quantification of flavivirus small RNAs in *M. persicae* and the Asian citrus psyllid (ACP) *Diaphorina citri* harboring the circulative plant pathogens PLRV and “Candidatus Liberibacter asiaticus” (*C*Las), respectively supports the hypothesis that a broad range of plant-infecting, circulative plant pathogens suppress hemipteran anti-viral immunity against ISVs.

## MATERIAL AND METHODS

### Total-RNA sequencing and data analysis

For virome analysis, parthenogenic colonies of *M. persicae* maintained on *N. benthamiana*, physalis (*P. floridana*), and turnip (*B. rapa*) were used. Each sample comprised groups of 30 aphids including four of each instar state, seven apterous and seven alates. Total RNA was extracted using cryogenic lysis and the Direct-zol^TM^ RNA MiniPrep kit (Zymo Reseach). Complementary DNA (cDNA) was synthesized using iScript Select Kit (Bio-Rad) as per the manufacturer’s instructions. RT-PCR assays targeting the mitochondrial cytochrome oxidase I (COI) gene from *M. persicae* were used to confirm cDNA integrity. Detection of the MpDNV was carried out to confirm endogenous viral infection in the aphid samples (Table S1). RNA samples were subjected to ribodepletion to remove the ribosomal RNA (rRNA) followed by cDNA library synthesis. HTS was carried out on an Illumina® NovaSeq 6000 system amplifying paired-end reads (2x100 bp) at the Genomic High-Throughput Sequencing Facility at the University of California, Irvine.

Raw reads were curated and assembled following the methods described by Larrea-Sarmiento (Larrea-Sarmiento et al. 2022). Briefly, trimming of raw reads was performed using Trimmomatic (Bolger et al. 2014). Quality reads were *de novo* assembled using Trinity v2.9.1 (Grabherr et al. 2011) and rnaSPADES V3.15.3 (Bushmanova et al. 2019). The resulting contigs were annotated by BLASTx searches of the NCBI virus sequence database as of September 2022. An iterative mapping approach to extend viral contig lengths was done using the Geneious mapper plugin implemented in Geneious version 2022.1.1 (Kearse et al. 2012). ORF presence in the viral contigs was evaluated using ORFfinder https://www.ncbi.nlm.nih.gov/orffinder/ (accessed on March 23^rd^, 2023) and conserved domains were annotated using the NCBI Conserved Domain search tool https://www.ncbi.nlm.nih.gov/Structure/cdd/wrpsb.cgi (accessed on March 23^rd^, 2023).

### dsRNA isolation and analysis

To validate the discovery of a new ISV infecting *M. persicae* and its active replication in the aphid, dsRNA extractions were carried out. Mixed developmental stages of aphids, including nymphs and alate or winged adults reared on physalis were bulk collected. Approximately 0.25 g of aphids was cryogenically lysed for dsRNA extraction using C6288 cellulose (Sigma) as previously detailed (Khankhum et al. 2017). Cryoground tissue (∼0.5 g) of Mexican lime infected with citrus tristeza virus (CTV, ∼19 kb) was also extracted and served as a control. Samples were separated by electrophoresis in 0.8 % agarose gels and visualized using the GelRed Nucleic Acid Stain (Sigma-Aldrich).

### Validation of transcriptomics data

Results from bioinformatic analyses were validated by RT-PCR assays in 20 μL reactions using Green GoTaq Master mix (Promega). Specific primers for the novel virus infecting *M. persicae* were designed using Primer3Plus v.3.3.0 (Untergasser et al. 2012) (Table S1). For the detection of MpDNV, literature primers were used (Pinheiro et al. 2019). PCR assays were carried out following conditions as described (Table S1). The genome of MpFV was amplified using seventeen overlapping primer sets, and products were directly sequenced using the Sanger method. Genome termini were characterized by rapid amplification of complementary ends on poly-A tailed dsRNAs that were generated using *E. coli* Poly(A) Polymerase (New England Biolabs), and 5’ and 3’ ends characterized as detailed previously (Navarro et al. 2018). Products were cloned into pGEM-T Easy (Promega), and five clones per amplicon were sequenced.

Cleavage sites of the polyprotein were predicted by multiple protein sequence alignment of the ORF from the new flavivirus with other members within the *Flaviviridae* family that have known cleavage sites. Moreover, SignalP (6.0 https://services.healthtech.dtu.dk/services/SignalP-6.0/) was used for the prediction of signal peptides and their cleavage sites. Secondary structures of the 5’ and 3’ UTRs were assessed using the Vienna package RNAfold tool version 2.6.0 plugin in Geneious Prime 2023.2.1. Turner 1999 and Turner 2004 algorithms were used with default parameters with the exception of 25°C temperature was used for the predictions.

### Phylogenetic analysis

A multiple protein sequence alignment of the RNA-dependent RNA polymerase (RdRp) of the new ISV and *Flaviviridae* homologs retrieved by BLASTP searches in March 2023 was performed. Specifically, the protein region between the 6758-7169 aa encoded by the long ORF was aligned with seventeen homologous sequences from related flaviviruses. Two distantly related homologs from the genus *Pestivirus* were included in the alignment as an outgroup. The maximum likelihood method and a gamma distribution were predicted as the best model for the 20 sequences aligned with a total of 286 positions. Maximum likelihood phylogenies were inferred using 1,000 bootstrap replicates, implemented in MEGA 11.0.13 (Tamura et al. 2021).

### Transovarial transmission

To evaluate the vertical transmission of the new flavivirus, *M. persicae* adult aphids from colonies reared on physalis were used. Single adults were collected and transferred to detached physalis leaves. The progeny was immediately collected from the posterior ends of their mothers while being born and transferred to new detached physalis leaves, with one nymph per leaf. After seven days, newly developed adult aphids were collected and flash frozen. Total RNA extraction and cDNA synthesis were conducted as described above. Detection of MpFV in aphids was carried out in RT-PCR assays using Green GoTaq Polymerase (Promega) and the MpFV-Hel primers with an expected PCR product of 361 bp (Table S1). The experiment was repeated three times independently.

### Artificial diet for the delivery of the new ISV to aphid species

To test whether the new flavivirus can infect other aphid species, we delivered the virus to other aphid species using artificial diets. Parthenogenetic clones of *Aphis gossypii* (cotton aphid) maintained on cotton and cucumber, *Schizaphis graminum-F* (greenbug aphid)*, Rhopalosiphum padi* (bird cherry-oat aphids), and *R. maidis* (corn leaf aphid) maintained on barley were used. For the new ISV homogenate, 0.2 g of *M. persicae* aphids exposed for 48h to PLRV-infected hairy nightshade (*Solanum sarrachoides)* plants were collected and cryoground. Then, aphid homogenate was resuspended in 3.4 mL of 40% sucrose: 1X PBS in a 1:1 ratio. Artificial diet delivery of MpFV was achieved by adding 270 µL of flavivirus suspension into a parafilm sache (Fig. S1a). Aphids from different species were placed in small ventilated dishes and allowed a 72-h acquisition access period (AAP). Then, aphids were transferred to their natural plant host (described above) for a 72-h period of gut clearing to remove flavivirus virions that may have been ingested but not acquired. Single aphid specimens were collected for RNA extraction and cDNA synthesis was performed as described above. RT-PCR assays were carried out using the MpFV-Hel and MpFV-RdRp primers (Table S1). PCR bands were excised and sequenced directly using the Sanger method. Detection of PLRV was conducted in RT-PCR assays as a control as detailed by Schiltz et al, 2022 (Schiltz et al. 2022).

### Flavivirus data mining

To evaluate whether the new flavivirus could also be found in other *M. persicae* colonies around the world, public sequence read archive (SRA) datasets were mined using the open-source cloud-computing infrastructure database, Serratus, for RdRp hallmarks specific to MpFV. Sequence from the conserved domain of the RdRp (MpFV-RdRP; 492 amino acids) was used for searches against the palmID Viral-RdRp analysis tool (Edgar et al. 2022). SRA accessions that displayed RdRp sequence homologies greater than 90% were retrieved and used for reference-guided assembly. Briefly, quality trimmed reads were mapped against the complete MpFV genome using the Geneious mapper, as detailed above. Sequence homology was analyzed using multiple sequence alignments at nucleotide and protein levels on Geneious.

### Small RNA (sRNA) sequencing and data analysis in aphids

Data generated and collected by Pinheiro et al. (2019) deposited on GenBank under the BioProject PRJN514359 was used for detection of small RNAs (sRNA) mapping to the flavivirus genome. Previously, sRNA sequencing was carried out from aphids feeding on infected potato plants and artificial diets with PLRV, potato plants infected with PVY, and a healthy potato control. sRNA raw data was trimmed using sRNA_clean.pl provided by VirusDetect (Zheng et al. 2017) to remove adapter sequence CTGTAGGCACCATCAAT as well as low quality and short (<15nt) reads. Cleaned reads were aligned to the MpFV genome using bbmap in perfect mode (https://sourceforge.net/projects/bbmap/). The distribution of sRNA sizes was visualized using ggplot2 in R Studio. A one-way fixed effects analysis of variance (ANOVA) was performed on normalized counts of 22mer *M. persicae* sRNA reads mapped to MpFV. Reads per kilobase per million (RPKM) mapped to MpFV were calculated for each biological replicate and Tukey’s test was ran to assign significantly different groups (*P*<0.05).

### Quantification of flavivirus titer in aphids

To assess whether PLRV can modulate the flavivirus titer in aphids, we implemented a droplet digital PCR (ddPCR) assay to quantify MpFV genome copies in aphids. A PLRV infectious clone (Franco-Lara et al. 1999) was used to inoculate one-month-old hairy nightshade plants. Parthenogenic colonies of *M. persicae* were maintained on physalis and nine-day aged-synchronized adult insects were used for this experiment. Synchronized aphids were caged and allowed a 3-day AAP on leaves from healthy NY-129 (Red Maria) potato plants, PLRV-infected hairy nightshade plants, and potato plants (Desiree) expressing the N-terminal portion of the PLRV readthrough domain (^N^RTD) (Schiltz et al. 2022). After AAP, aphids were transferred to detached turnip leaves for a 3-day period of gut clearance (Fig. S1b). Single aphids were collected, resulting in 18 to 20 biological replicates per treatment. Total RNA was extracted using the Direct-zol^TM^ RNA MiniPrep kit (Zymo Reseach), including an on-column DNase digestion treatment as per the manufacturer’s instructions. About 88.88 ng/µL of total RNA was added to the cDNA synthesis reactions using the iScript Select Kit (Bio-Rad).

For semi-quantitative PCR, 2 μL of cDNA was added to the RT-PCR reaction mixture along with 10 μL of 2x GreenTaq polymerase (Promega, Madison), 1 μL of each MpFV-Hel forward and reverse primers (10 μM), and 6 μL of nuclease-free water. To measure the titer of MpFV, we implemented an RT-ddPCR assay in the QX200 ddPCR system (Bio-Rad). Briefly, 2 μL of normalized aphid cDNA was added to the ddPCR reaction mixture along with 10 μL of 2x EvaGreen SuperMix, 7 μL of nuclease-free water, and 0.5 μL of each of the following specific primers to the MpFV RdRp coding region: FP 5′-TCATCGAAGGCCTATGTCGC-3′ and RP 5′-TCAGTATCTCGCCTGACCCC-3′. Droplets were generated using the QX100 droplet generator (Bio-Rad) using 70 μL of droplet generator oil and 20 µL of reaction. Water-in-oil droplets (∼42 μL) were transferred to a 96-well plate, and PCR was carried out as follows: 95°C for 5 min, followed by 40 cycles of 94°C for 30 sec, and 60°C for 60 sec, followed by 4°C for 5 min and 90°C for 5 min for signal stabilization. After PCR, reactions were read on the QX200 ddPCR machine using the FAM channel. Droplet counts were analyzed using QuantaSoft v.7.1 (Bio-Rad). The experiment was repeated twice independently. Data was analyzed and visualized using ggplot2 in R Studio.

### Correlation analysis

To test whether there was a correlation between the titers of PLRV and the new flavivirus in aphids, the titer of both viruses was measured from the same individual aphid samples. Twenty-two samples were randomly selected from the assays described above, and PLRV titer was measured by ddPCR as described by (Schiltz et al. 2022). The Spearman and Kendall correlation methods were used to measure a correlation coefficient and *P*-values.

### Transient in planta delivery of viral suppressors

To evaluate the effect of the viral gene silencing suppressors on flavivirus titer, aphids were exposed to *N. benthamiana* transiently expressing proteins as follows: two mL of *Agrobacterium tumefaciens* suspensions (O.D. = 0.5) expressing PLRV P0, tomato bushy stunt virus (TBSV)-p19 (p19), and the PLRV infectious clone were infiltrated on four leaves per plant, three plants per construct. At three days post infiltration (dpi), nine-day-old apterous *M. persicae* adult aphids were transferred (three aphids/leaf, four leaves/plant, three plants/treatment) and caged on the construct-expressing leaves as well as un-infiltrated (control) leaves for 48 h. Then, three aphids feeding in different leaves were randomly collected as one sample with a total of 6 samples per treatment (Fig. S1c). Total RNA extraction, DNase treatment, cDNA synthesis, and quantification of the flavivirus by RT-ddPCR were carried out as detailed above.

### Flavivirus tissue tropism determination

To evaluate the tissue tropism of the new flavivirus, we performed aphid dissections. *M. persicae* colonies established on physalis were used in this experiment. For the control group, third-instar aphids were transferred to healthy Red Maria potato plants and allowed an AAP of seven days. For aphids feeding on PLRV-infected potato plants (Red Maria), an AAP of five days was used followed by a gut clearing period of two days on healthy potatoes. Aphids were collected in groups of three and samples were stored at −80°C. Under the dissecting scope, frozen aphids were used in dry dissections to avoid cross-contamination in an arena sitting on ice: a plastic plate containing the paper side of parafilm. The head (brain and salivary glands) along with the prothorax first thoracic segment bearing two legs were separated from the rest of the body (mesothorax and/or abdominal segment) and treated as two separate but paired samples. Each sample corresponded to a pool of three heads or bodies. Flame-sterilized tools were used in between aphid samples to eliminate contamination. Total RNA extraction, DNase treatment, cDNA synthesis, and quantification of the new flavivirus by RT-ddPCR were carried out, as detailed above.

### Evaluation of flavivirus replication in plants

To assess if the new flavivirus infecting *M. persicae* is detectable in plants infested with *M. persicae*, *N. benthamiana* plants were agro-inoculated with binary vectors expressing the green fluorescent protein (GFP) and the PLRV infectious clone. These plants and an uninfiltrated control plants were infested with 10 *M. persicae* adults in a clip cage to confine aphid feeding to a single site, which we hypothesized would maximize the local concentration of the flavivirus in the plant. After three days, the aphids were removed, and three plant punches were taken at the site of inoculation and infestation. Three more punches were taken from the newest developing leaves from each treatment and control to test for systemic movement of the new flavivirus. The plant punches were flash-frozen, and RNA was extracted using the Qiagen RNeasy Mini kit following manufacturer’s instructions. RT-PCR detection of MpFV was performed as detailed above.

### Small RNA sequencing and data analysis in psyllids

To investigate *D. citri* antiviral immunity in the presence and absence of *C*Las, small RNAs were sequenced from *C*Las(+/-) adult and nymph ACP. A total of 40 insects were pooled into bio-replicates for sRNA extraction and sequencing for the four groups, where both *C*Las-infected datasets contained 5 bio-replicates and healthy datasets contained 4 bio-replicates. Raw sRNA sequencing data were filtered for adapter sequence TGGAATTCTCGGGTGCC as well as low quality and short (<11nt) reads using sRNA_clean.pl provided by VirusDetect (Zheng et al. 2017). Cleaned sRNA reads were mapped to the *Diaphorina citri* Flavi-like virus genome (NC_030453.1) using bbmap (https://sourceforge.net/projects/bbmap/) in perfect mode allowing no mismatches. To investigate the size distribution of sRNAs mapping to the DcFLV genome percentages of mapped reads were calculated by dividing length specific read counts by the total cleaned reads of each respective replicate. The percentages for mapped sRNAs of each length were then averaged across biological replicates and plotted. A one-way fixed effects analysis of variance (ANOVA) was performed on normalized counts of 21mer *D. citri* sRNA reads mapped to DcFLV. Reads per kilobase per million (RPKM) mapped to DcFLV were calculated for each biological replicate and Tukey’s test was ran to assign significantly different groups (*P*<0.05). Statistics were visualized with ggplot2 in R studio.

### MiSiPi Analysis

Short interfering RNAs were specifically investigated using MiSiPi (https://www.ncbi.nlm.nih.gov/pmc/articles/PMC10197562/) after 21mers and 22mers were observed to dominate both the *M. persicae* and *D. citri* flavivirus sRNA profiles. Sorted and indexed bam files of each replicate, produced from mapping sRNA reads to MpFV and DcFLV, were combined using samtools merge and fed into MiSiPi’s siRNA_function. Heatmaps generated depict the size distribution of duplexed siRNAs containing Dicer-2 3’ 2nt overhangs.

## RESULTS

### A novel flavivirus infecting the green peach aphid

Annotated contigs from the *M. persicae* HTS dataset revealed sequence similarity to the already reported DNA virus infecting *M. persicae*, MpDNV, and also to members of the *Flaviviridae* family. Protein identities below 58% with other flavivirids suggested the presence of a new flavivirus infecting the green peach aphid. The closest relative is Macrosiphum euphorbiae virus 1 (22,780 nts, NC028137). The complete genome sequence for the new flavivirus is 23,226 nts (accession XXXXXX) and is predicted to encode a long ORF of around ∼22.5 kb (315 to 22,790). Presence of the new ISV in the green peach aphid was validated *in vitro* using dsRNA extractions from our *M. persicae* colonies. A clear band between 20 to 25 kb was observed on electrophoresis gels from the dsRNA *M. persicae* sample, while a band of the expected size, ∼19 kb, was observed for the dsRNA extraction control CTV-plant tissue (Fig. 1a), corresponding to the size of the CTV genome. Moreover, RT-PCR assays using specific primers targeting two regions corresponding to the RdRp and Hel genes of the MpFV genome confirmed the infection of the *M. persicae* colonies from three plant hosts: *N. benthamiana*, physalis and turnip. No bands were obtained when testing other aphid species maintained in the same room by our group (Fig. S2). These results corroborate the infection of *M. persicae* by a new flavivirus. We propose the name of Myzus persicae flavivirus (MpFV) for this novel ISV infecting the polyphagous green peach aphid.

**Fig. 1.**
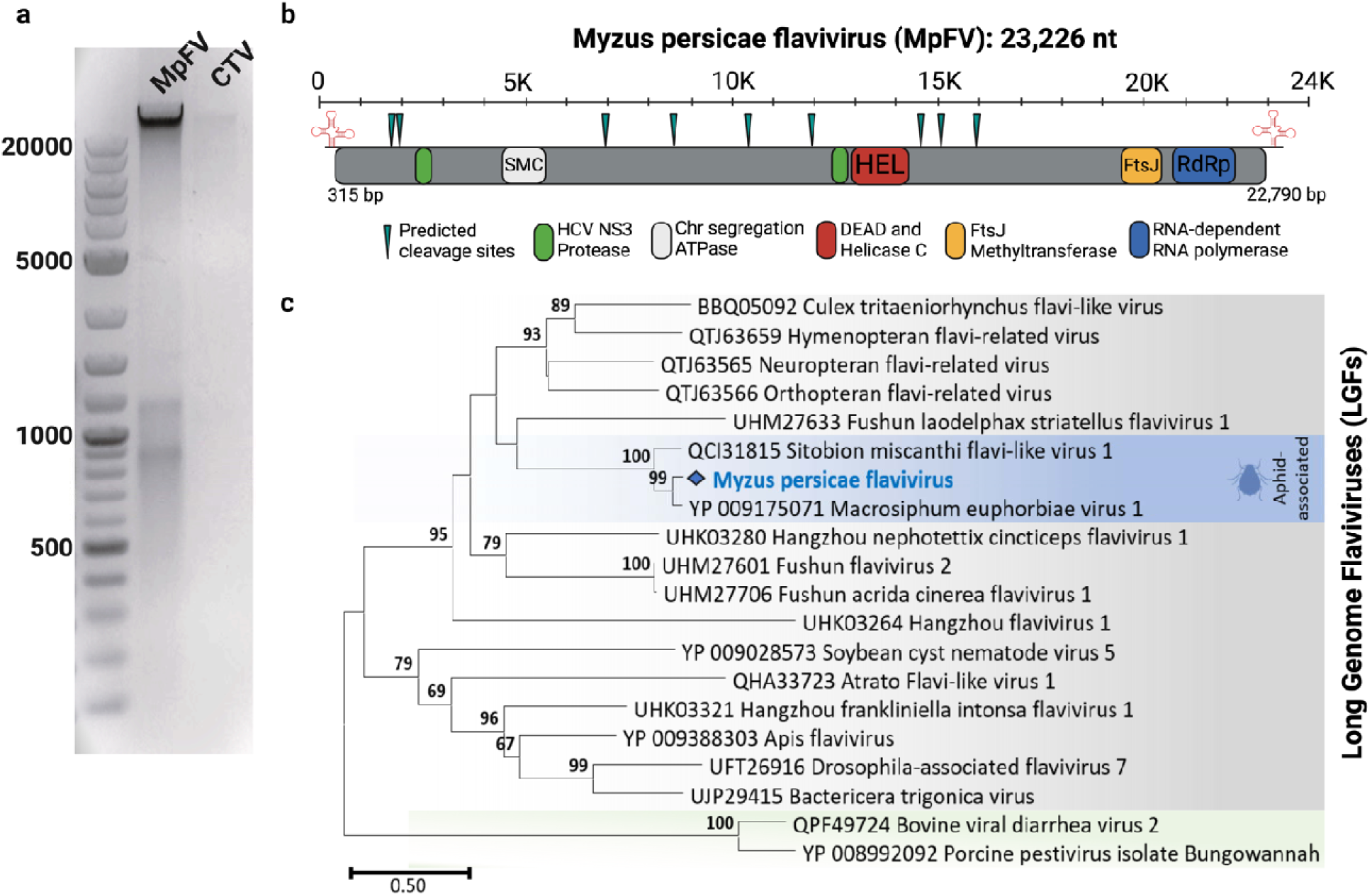
Schematic representation of the Myzus persicae flavivirus (MpFV) genome and phylogeny. **(a)** dsRNAs extractions from citrus infected with citrus tristeza virus (CTV, ∼19 kb) and from *M. persicae* infected with MpFV (∼23 kb) run in 0.8% agarose gel electrophoresis. A 1 Kb-plus DNA Ladder (ThermoFisher Scientific) is in the left lane with its corresponding prominent DNA bands with different lengths. **(b)** Open reading frame (ORF) of MpFV with the predicted conserved domains flanked by the 5 and 3 untranslated regions. Predicted cleavage sites are represented on green. **(c)** Phylogenetic analysis of MpFV using the identified, conserved domain from the RNA-zdependent RNA polymerase (RdRp) protein sequence (aa 6758 to 7169). MpFV was grouped within the LGF cluster. Two virus species from the Pestivirus genus (dual-host) were used as an outgroup.

### Genome organization and phylogeny of Myzus persicae flavivirus (MpFV)

Pairwise alignment of the flavivirus contigs recovered from the three datasets representing *M. persicae* in different hosts showed a 99.9% nucleotide identity among one another. *In silico* analysis using NCBI conserved Domain search tool (https://www.ncbi.nlm.nih.gov/Structure/cdd/wrpsb.cgi; accessed on March 2023) showed presence of conserved domains from helicases, NS3-proteases, Fts-J methyltransferase, and RdRp, typical of other LGFs of the *Flaviviridae* family. Analysis of cleavage sites from MpFV supports the hypothesis that the viral genome is translated as a polyprotein and cleaved by virus-encoded and/or host proteases to produce 10 mature proteins. Based on the genome organization of the *Flaviviridae* family, the long ORF of MpFV is expected to encode 10 mature proteins starting at the 5 with the three structural proteins capsid (C), precursor-membrane (M) prM, and envelop (E), followed by the seven non-structural proteins (NS): NS1-NS2A-NS2B-NS3-NS4A-NS4B-NS5 (Fig. 1b). The MpFV viral genome encoding the polyprotein is likely flanked by 5 capped and 3 non-poly-A tailed UTRs with 314 and 395 nucleotides in length, respectively. Stem-loop secondary structures were observed in both non-coding regions (Fig. S3).

Phylogenetic analysis using the protein sequence from the identified RdRp conserved domain revealed that the MpFV is a new member of the *Flaviviridae* family clustered in the unclassified group of LGF. As expected, all LGF are distinct from pestiviruses, Bovine viral diarrhea virus 2 and Porcine pestivirus, both classified as dual-host viruses since infect vertebrates and invertebrates. MpFV is closely related to two other ISV (99-100% bootstrap support), Sitobion miscanthi flavi-like virus 1 and Macrosiphum euphorbiae virus 1, characterized to be aphid-associated viruses (Fig. 1c).

### Vertical and horizontal transmission of MpFV

We tested whether MpFV is vertically transmitted from parents to the offspring. Three *M. persicae* generations were tested: parents (F1), offspring-1 (F2), and offspring-2 (F3). Nymphs were collected at the moment of birth from parent females to discard any possible honey-dew horizontal transmission. A transmission of 100% was obtained when individual aphids from the offspring were tested by RT-PCR assays using the primer set targeting the MpFV-polymerase. Four aphids died before collection, one from the generation F1 and three from the generation F3, thus these were not tested.

Moreover, we also tested whether MpFV can be horizontally transmitted to the other aphid species by artificial diet with MpFV-infected *M. persicase* homogenate and by evaluating the presence of MpFV in plant tissue after aphid infestation. For the aphid experiments, RT-PCR assays targeting the polymerase (MpFV-RdRp) and the helicase (MpFV-Hel) regions were used to assess transmission of MpFV. The number of positive aphids was different when using oligonucleotides targeting the RdRp and/or Hel (Table S2). A total of 176 aphids were tested from five different species with an overall transmission efficiency of 12.9% and 14% for the RdRp and Hel, respectively (Table 1). *R. maidis* aphids were found dead before the gut clearing period in replicates 1 and 2, suggesting that *M. persicae*-MpFV homogenate was toxic for the corn leaf aphid. Although, some *R. maidis* individuals survived after the gut clearing period, none acquired MpFV. The greenbug aphid, *Schizaphis graminum*-F, was the most efficient aphid species to acquire MpFV with transmission rates of 18% and 12%. The PCR products were sent to Sanger sequencing and submitted to NCBI (Table S2). All sequences showed 100% homology with *M. persicae* sequence with the exception of one sample, of *A. gossipii* from cucumber, from replicate 2 that presented a single point mutation (C121T). For the plant tissue, the RdRp primers in RT-PCR assays were used, and no MpFV was detected in any of the aphid-infested plant samples evaluated (data not shown).

**Table 1.** Horizontal transmission of Myzus persicae flavivirus (MpFV) to several aphid species by artificial diet feeding.

|  |  |  | MpFV-polymerase (RdRp) |  |  | MpFV-Helicase (Hel) |  |  |
| --- | --- | --- | --- | --- | --- | --- | --- | --- |
| Aphid species |  | total | RT-PCR Positive |  | Transmission (%) | RT-PCR Positive |  | Transmission (%) |
| After AAP* | *R.padi* | 26 | 6 | 6/26 | 23 | 3 | 3/26 | 12 |
|  | *R. maidis* | 25 | 2 | 2/25 | 8 | 8 | 8/25 | 32 |
|  | *A. gossypii*-cucumber | 11 | 1 | 1/11 | 9 | 1 | 1/11 | 9 |
|  | *A. gossypii*-cotton | 19 | 2 | 2/19 | 11 | 1 | 1/19 | 5 |
|  | *S. graminum* | 16 | 3 | 3/16 | 19 | 5 | 5/16 | 31 |
| After gut clearing† | *R.padi* | 8 | 0 | 0 | 0 | 0 | 0/8 | 0 |
|  | *R. maidis* | 0 | 0 | - | - | - | - | - |
|  | *A. gossipy*-cucumber | 33 | 2 | 2/33 | 6 | 4 | 4/33 | 12 |
|  | *A. gossypii*-cotton | 23 | 4 | 4/23 | 17 | 1 | 1/23 | 4 |
|  | *S. graminum* | 17 | 3 | 3/17 | 18 | 2 | 2/17 | 12 |
| Total aphids tested |  | 178 | 23 | 23/178 | 12.9 | 25 | 25/178 | 14 |

\* Dead aphids collected before gut clearing period
† Aphids were transferred from artificial diets to their natural host for a 72-hour gut-
clearing period. *Rhopalosiphum maidis*, *R. padi*, and *Schizaphis graminum* were moved
to barley; *Aphis gossypii*-cucumber to cucumber, and *A. gossypii*-cotton to cotton.

### MpFV genetic diversity and insights into aphid tissue tropism

Data-driven virus discovery was used to explore the presence and natural diversity of MpFV in aphid sequencing datasets deposited by researchers world-wide. The conserved domain and the hallmarks of the RdRp sequence in MpFV was used for searches against the palmID Viral-RdRp analysis tool on the Serratus database. Biosample accessions showing identities >90% were used to retrieve datasets from the SRA database. Analyzes of the geographic distribution showed a worldwide presence of MpFV sequences in *M. persicae* datasets from China (SAMN07709675, 2016), Italy (SAMN12860378, 1996), England (SAMN08668666, 2014), Germany (SAMN12860380, 1967), Scotland (SAMEA3505165, 2015); and United States (Texas, SAMN04867904). Surprisingly, MpFV sequences were also detected in a dataset derived from a native central and southern Europe plant collected from Germany (SAMN15369604), *Blitum bonus-henricus*, commonly known as Good-King-Henry (Fig. S4). Normalized data used for nucleotide and protein alignments showed identities over 91% and 97%, respectively (Fig. S4). These findings highlight not only a global distribution of MpFV but also its genome sequence conservation.

The Scotland samples were comprised of decapitated aphid bodies and aphid heads. Thus, normalized data from the Scotland sample was used to develop hypotheses about MpFV tissue tropism. Normalized reads were mapped against the body (ERR983180-82) and head (ERR983177-79) datasets with a p-value of 0.0103. These data sets contained a higher number of MpFV reads in the body tissues as compared to the head samples, suggesting that MpFV has a tissue tropism for tissues within the body rather than the head (Fig. 2b). An evaluation of MpFV titer from excised heads and bodies of PLRV-viruliferous *M. persicae* from our lab colony showed the same tissue tropism as the Scotland samples (Fig. 2a), providing further support for MpFV tissue tropism within tissues of the aphid metathorax or abdomen.

**Fig. 2.**
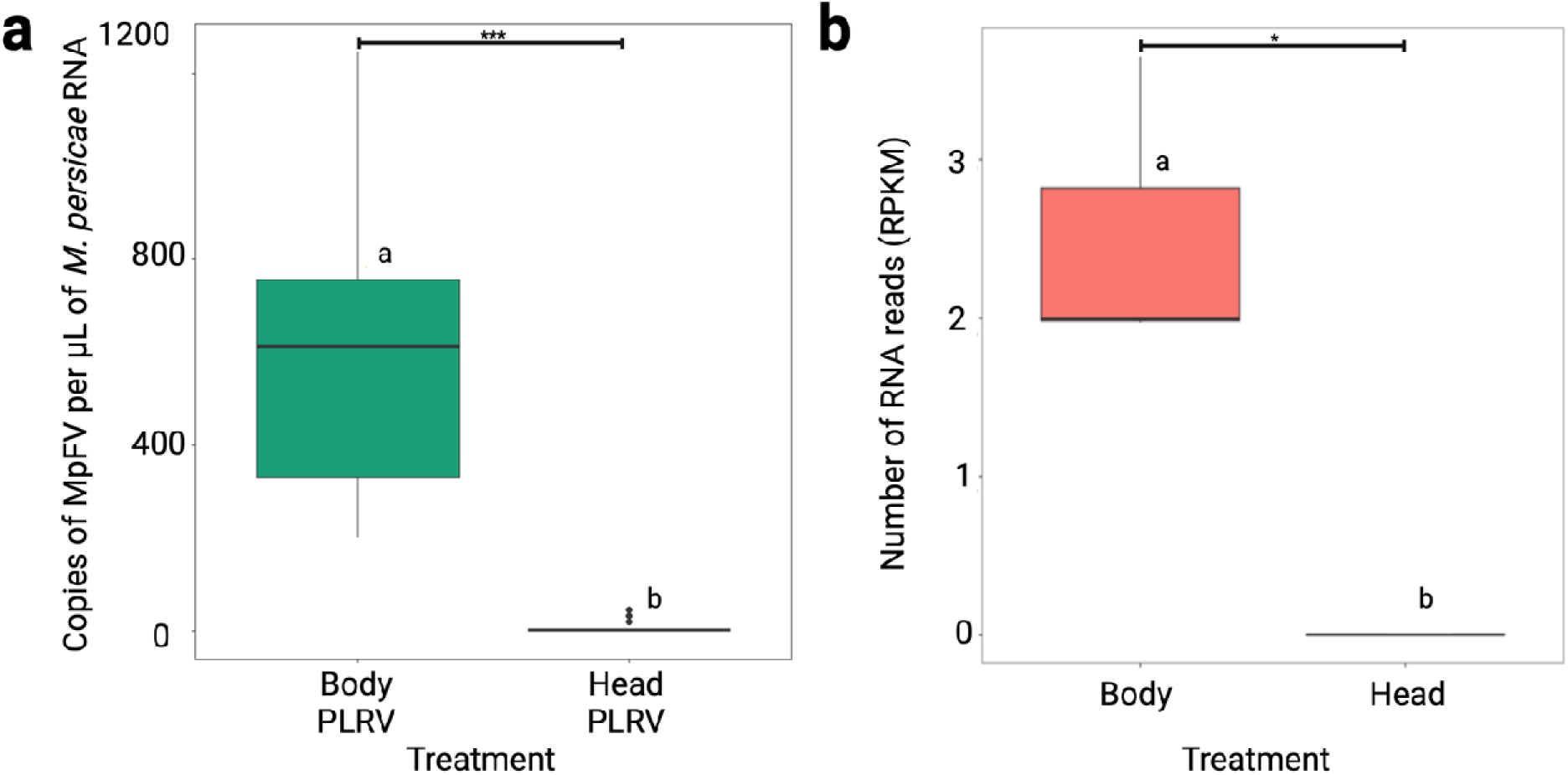
Tissue tropism of Myzus persicae Flavivirus (MpFV) within *M. persicae*. **(a)** Absolute quantification of MpFV showing infection within the aphid body (metathorax and/or abdominal segment) rather than the heads (brain and/or salivary glands) in PLRV-viruliferous aphids. ***Turkey test *P*-value = 0. **(b)** Tissue tropism of MpFV in the *M. persicae* Scotland sample. Normalized reads (reads per kilobase per million) were mapped against the body (ERR983180-82) and head (ERR983177-79) with a \**P*-value of 0.0103.

### PLRV modulates the aphid’s immune response to MpFV

Previous work from our group showed that PLRV interferes with the aphid’s immune response to the ISV, MpDNV in *M. persicae*, resulting in higher titers of MpDNV in PLRV viruliferous aphids (Pinheiro et al. 2019). To assess whether PLRV also modulates the *M. persicae* immune response against MpFV, we first re-analyzed the sRNA datasets from Pinheiro et al. 2019 for reads matching to MpFV. Profile analysis of MpFV from PLRV-viruliferous aphids were compared to non-viruliferous aphids. Aphid libraries contained sRNA reads matching to MpFV varying from 15 to 40 nt in length. An abundant production of 15-22 nt size of siRNA with less production of >23 nt size was observed when aphids fed on healthy plants, sucrose diet, and PVY-infected plants. In contrast, no sRNA reads matching to MpFV were detected in PLRV-viruliferous aphids, regardless of whether the aphids acquired PLRV from artificial diets or infected plants (Fig. 3a), showing PLRV acquisition interfered with the production of sRNAs from the flavivirus genome during virus replication (Fig. S5).

**Fig. 3.**
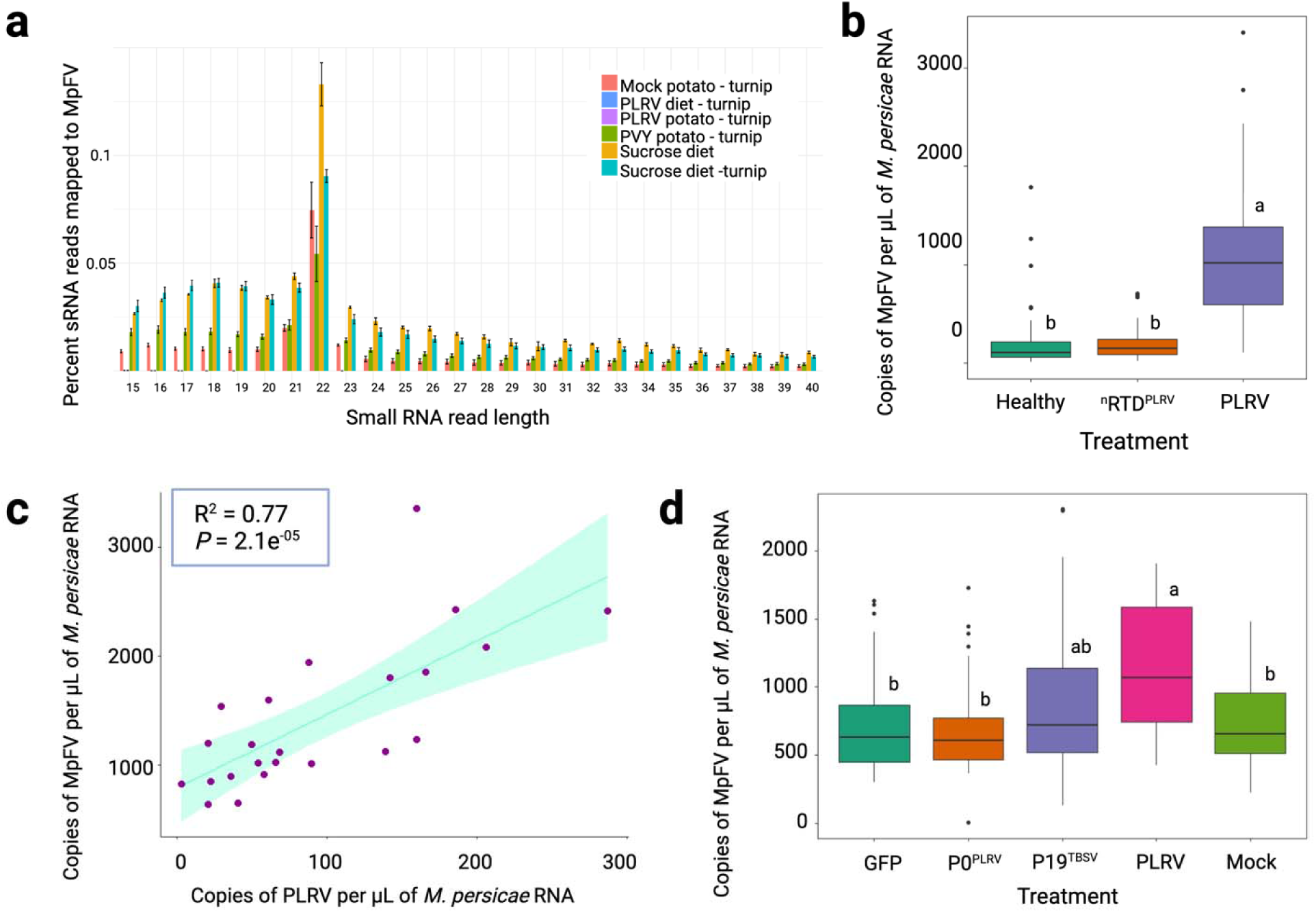
Regulation on *Myzus persicae* immune response and modulation of Myzus persicae flavivirus (MpFV) titer by potato leafroll virus (PLRV). **(a)** Distribution of small RNA (sRNA) reads generated in *M. persicae* aligned to the genome of the new insect specific virus (ISV), MpFV. Aphids were given a 72-h acquisition access period (AAP) on the different treatments followed by a gut clearing period of 72-h. ANOVA revealed a significant difference in 22mer reads mapped to MpFV among conditions with a p-value of 2.54e-07. **(b)** Absolute quantification of MpFV in individual *M. persicae* aphids by droplet digital PCR (ddPCR). Aphids were given the same AAP and gut clearing periods as described above. **(c)** Correlation of PLRV and MpFV titers. Spearman (R= 0.77, *P*=2.1e-05) and Kendall (R=0.59, *P*=7.4e-05) methods revealed a positive association between the titer of PLRV and MpFV within aphids. **(d)** Quantification of MpFV on aphids (3 adults/biorep) that fed on *Nicotiana benthamiana* plants transiently expressing the PLRV infectious clone, PLRV P0, tomato bushy top virus P19, the green fluorescent protein (GFP) or mock-infiltrated plants.

To further evaluate the relationship between MpFV and PLRV titer, we co-measured MpFV and PLRV titers using ddPCR on individual aphids that had acquired PLRV. These experiments revealed that PLRV acquisition by aphids increases the titer of MpFV; however, the increase in MpFV titer is not mediated by the sole presence of the PLRV minor structural protein, the N-RTD, *in planta* (Fig. 3b, *P*-value = 0.0 using a Tukey test). MpFV concentration on individual aphids that fed on PLRV-infected plant tissue was almost three times higher than those that fed on non-infected plants (Fig. 3B). A correlation analysis between PLRV with MpFV titer in individual aphids using Spearman (R^2^= 0.77, *P*=2.1e-05) and Kendall (R^2^=0.59, *P*=7.4e-05) methods reveals a positive association between the titers of PLRV and MpFV (Fig. 3c), which was unexpected because although PLRV is a circulative plant pathogen, the virus does not replicate in aphids (Pinheiro et al. 2019).

Next, we tested whether the increase in MpFV titer mediated by PLRV was mediated by the PLRV silencing suppressor protein, P0, as previously reported for MpDNV (Pinheiro et al. 2019). Assays using *N. benthamiana* transiently expressing different constructs were carried out to evaluate this hypothesis: P0, the viral suppressor protein, p19 from tomato bushy top virus, GFP and the whole PLRV virus. Absolute quantification of MpFV from the aphids collected revealed that PLRV P0 alone does not induce a titer change of MpFV; however, p19 triggered an intermediate titer increase between the PLRV and the control treatments (Fig. 3d).

### *C*Las regulates the *D. citri* response against Diaphorina citri flavi-like virus (DcFLV)

To test the hypothesis whether modulation of the anti-viral immune system of hemipteran vectors by circulative pathogens is unique to *M. persicae* and PLRV, we drew upon an existing dataset in our lab derived from *D. citri,* a hemipteran insect vector of the circulative, propagative citrus-infecting pathogen, *C*Las (Hosseinzadeh et al, in prep.). Multiple biological replicates from *C*Las + and *C*Las – adult and nymph *D. citri* reared on either healthy or *C*Las (+) citron were pooled for small RNA isolation and sequencing. Small RNA reads were mapped to the DcFLV genome and the lengths of the matches plotted to examine the *D. citri* anti-viral immune response (Fig. 4). *C*Las (-) nymphs produced the most abundant small RNAs, followed by *C*Las (-) adults. Similar to PLRV-viruliferous aphids, *C*Las (+) adults and nymphs produced almost no sRNA reads matching to DcFLV (Fig. 4b), showing that *C*Las (or a biomolecule produced or induced by *C*Las in the infected citrus plant) suppresses anti-viral immunity against ISVs in *D. citri*. The most abundant size of sRNA produced in aphids against MpFV was 22 nt while in psyllids against DcFLV, it is 21 nt (Fig. 3a, Fig. 4a, Fig. S6).

**Fig. 4.**
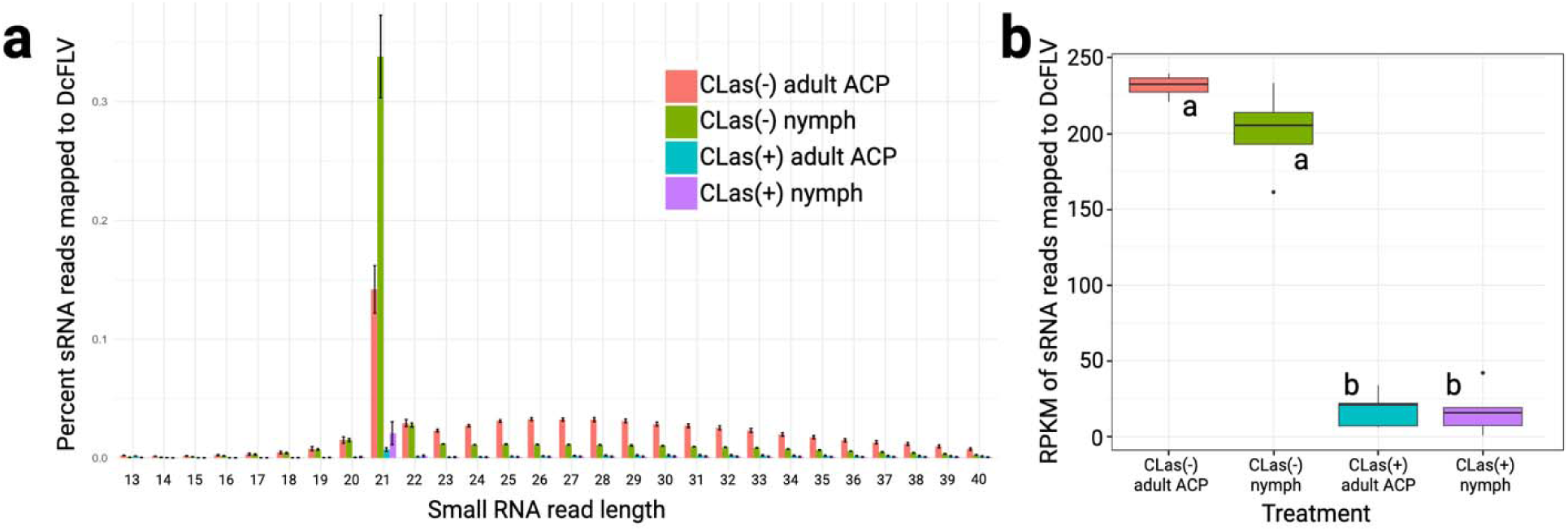
Small RNA analysis of the *Diaphorina citri* flavi-like virus (DcFLV) genome in adult and nymph *D. citri* reared on *C*Las (+) or (-) citron plants. **(a)** Size distribution of the reads mapped to DcFLV in *C*Las (+) and (-) nymph and adult insects. ANOVA revealed a significant difference in 21mer reads mapped to DcFLV among conditions with a p-value of 9.32e-08. **(b)** Reads per kilobase per million (RPKM) mapped to DcFLV genome from different *D. citri* small RNA datasets. ANOVA revealed significant differences in RPKM values with a *P*-value of 1.48e-11. Tukey’s test determined two significantly different groups (a & b). Nymphs and adults were collected from the same colonies for each biological replicate.

## DISCUSSION

We report the discovery of a new ISV infecting *M. persicae*, MpFV. The presence of MpFV infecting *M. persicae* was confirmed using RT-PCR assays and dsRNA extraction from aphid homogenate (Fig. 1A). The genome of MpFV is characterized by a long single-stranded RNA molecule (>20 kb) with stem-loop secondary structures on both ends. In other flaviviruses, these structures are involved in the initiation of RNA synthesis and virus replication, generation of subgenomic flavivirus RNAs and evasion of host immune response (Ng et al. 2017). Similar to other LGF and members of the family *Flaviviridae (Paraskevopoulou et al. 2021)*, our data predicts that MpFV codes for a single polyprotein that is cleaved into 10 mature proteins involved in the virus replication and infection cycles within its aphid host. This new flavivirus is related to other aphid-associated flaviviruses within the unclassified cluster of LGF of the family *Flaviviridae*.

Vertical transmission of MpFV was observed with 100% efficiency. Vertical transmission of other arboviruses that are members within the *Flaviviridae* family has been previously reported (Cook et al. 2006; Saiyasombat et al. 2011; Sang et al. 2003), including DcFLV in *D. citri* (Rashidi et al. 2022). Acquisition of MpFV by aphid species different from *M. persicae* shows that horizontal transmission via oral ingestion between aphid species is feasible with a 12-14% transmission efficiency. These results are consistent with a previous study that found that ISVs are efficiently transmitted via a vertical and/or sexual routes (Bolling et al. 2012). In contrast, horizontal transmission for members within the *Flavivirus* genus has been observed within the dual-host flaviviruses (DHV) group, characterized to infect arthropods and horizontally transmitted by feeding to vertebrate hosts (Blitvich and Firth 2015; Kuno 2007; Vasilakis and Tesh 2015). As other ISVs, MpFV is expected to have a defined host range. We did not detect MpFV in *N. benthamiana* cells, suggesting that MpFV cannot be transmitted through plants, although we cannot unambiguously state this as only one species of plants was evaluated in our experiments. Certain ISVs can be detected in plant RNA samples (Rashidi et al. 2022), horizontally transmitted to their herbivorous insect hosts via plants or plant surfaces contaminated with honeydew (Jia et al. 2020; Lu et al. 2020; Murakami et al. 2014). These attributes will need to be further explored to understand whether MpFV can be used as a biocontrol agent similar to the bacteria *Wolbachia* (Nasar et al. 2015; Öhlund et al. 2019).

Data mining using the Serratus database (Edgar et al. 2022) allowed us to explore the genetic diversity and geographic distribution of the new ISV characterized in our study. MpFV presented a worldwide distribution, with its presence inferred from North America (USA), Europe (Italy, England, Germany, and Scotland), and Asia (China). Interestingly, the dataset from Germany did not derive from *M. persicae* and instead originated from *Blitum bonus*-*henricus*, a plant native to Europe (Fig. S4). We cannot rule out the possibility that the plant sample used to generate the SRA dataset was free of insects possibly harboring MpFV, which could have included the polyphagous *M. persicae*. Moreover, the genetic diversity observed in different MpFV isolates from several countries did not surpass 91% nucleotide divergence, which translated to a more conserved protein identity level of over 97% (Fig. S4). RNA viruses with divergent genomes but conserved proteins have been previously reported to infect different hosts (Charon et al. 2022). Data mining also allowed us to generate new biological hypotheses. Specific datasets from decapitated aphid bodies and heads suggested that MpFV accumulates in bodies rather than in heads (Fig. 2B). To corroborate this tissue tropism, insect dissections and absolute quantification using ddPCR assays demonstrated that MpFV also accumulates in *M. persicae* abdomens rather than in heads of the insects from our colonies (Fig. 2A). Flaviviruses have been shown to infect and replicate in different insect tissues; however, some viruses may present specific host-virus interactions and may limit their replication to certain body tissues such as abdominal and thoracic fat bodies (Huang et al. 2014).

The suppression of aphid antiviral immunity by PLRV against MpFV (this study) and MpDNV (Pinheiro et al. 2019) may be related to the vector manipulation hypothesis, where plant viruses enhance their spread to healthy new hosts through their effects on mobile insect vectors (Ingwell et al. 2012). Data collected in the present study demonstrated the ability of the aphid-borne circulative-transmitted plant virus, PLRV to manipulate the immune response of the vector, *M. persicae*, as well as its ability to modulate the titer of MpFV within the insect. This trend was not observed for the aphid-borne non-persistent-transmitted plant virus, PVY (Fig. 3A). For MpDNV, PLRV-viruliferous aphids demonstrated a shift in the production of longer sRNAs against MpDNV when PLRV was acquired from infected plants. In contrast, no small RNAs were produced by PLRV-viruliferous aphids derived from MpFV when PLRV was acquired from either infected plants or artificial diet. The PLRV P0 silencing suppressor protein was found to mediate the interactions between PLRV and MpDNV via expression in the plant, either directly or indirectly, but this was not the case for MpFV. Together, these data support the hypothesis that there are at least two distinct molecular mechanisms used by PLRV to modulate the aphid anti-viral immunity, one dependent on P0 via the plant and one dependent on the virion or plant-virion protein/macromolecular complexes, which may not be dependent on the partial region of the PLRV NRTD used in our experiments. PLRV acquisition has also been reported to reduce the titer of the aphid’s primary obligate bacterial symbiont *Buchnera aphidicola* (Patton et al. 2021), but whether the ISVs and *Buchnera* are responding to similar molecular changes in the aphid as a result of PLRV acquisition is not known. The positive correlation between the titers between PLRV and MpFV within *M. persicae* provides evidence for the vector manipulation hypothesis at the molecular level. More research is needed to understand whether MpFV regulates PLRV transmission directly. Positive associations between co-infecting flaviviruses have been described in mosquitos. Of interest, Culex flavivirus, an ISV globally distributed in mosquitoes, occurs in co-infection with West Nile virus (Newman et al. 2011). These vector-ISV-viral pathogen interactions demonstrate that viruses with distinct transmission modes modulate insect immunity to benefit the transmission of both viruses, which is critical to understand at an ecological and evolutionary level due to the impacts on plant and animal health.

*C*Las is a Gram-negative, fastidious bacterium associated with Huanglongbing (HLB, also referred to as citrus greening disease) in the United States, Asia and elsewhere. A previous report by Rashidi and colleagues (2022) showed a positive association between *C*Las and DcFLV in field collected psyllids, similar to the positive association we found with PLRV and MpFV titer here as well as our previous report of higher MpDNV titers in PLRV viruliferous aphids. Thus, we tested whether the anti-viral immune response of *D. citri* to the DcFLV was changed when adult or nymph psyllids were bacteriliferous for *C*Las. Production of siRNAs indicates active viral replication and the corresponding anti-viral immune response. In the *D. citri* colonies we examined, we measured the highest number of reads in nymph, *C*Las (-) *D. citri*, which is consistent with the report of Rashidi et al. (2022) showing higher DcFLV titers in *C*Las (-) nymphs as compared to adults. Strikingly, almost no DcFLV reads were detected in *C*Las (+) adult or nymph *D. citri*, mirroring the findings we report for MpFV in *M. persicae.* These results point to an evolutionary conserved mechanism for circulative plant pathogens in different genera (viruses and bacteria) to modulate the anti-viral immune response in their hemipteran vectors. The major species is sRNA in aphids and psyllids produced against their flaviviruses is 22-nt and 21-nt respectively, supporting the hypothesis of different functionality in dicer enzymes, distinct dicing substrates, and/or the impact of length on sRNA efficacy in these different insect hosts, such as seen in *Drosophila melanogaster* and mosquitoes (Sabin et al. 2013). We do not yet know the mechanism for how *C*Las would influence *D. citri* anti-viral immunity, but we speculate it may be due to the expression of *C*Las effector proteins during *C*Las replication in the insect or the plant. Bacterial effector proteins help the pathogen evade the host immune response, determine tissue tropism, among other functions (Huang et al. 2020). A number of different *C*Las effector proteins have already been characterized to be expressed in citrus, psyllids or both organisms (Yan et al. 2013).

Transmission of plant pathogens by hemipteran insects is broadly defined based on the length of time the pathogen remains associated with the vector (Gray et al. 2014). We previously showed that non-persistent pathogens, such as Potato virus Y (PVY), which are not acquired into the aphid body, do not induce changes in the aphid’s antiviral immune system pathways (Pinheiro et al. 2019). In contrast, the similarities between PLRV, a viral pathogen and *C*Las, a bacterial pathogen, in their ability to modulate antiviral immunity in two insect vector species within in different superfamilies in the Hempitera suborder Sternorrhyncha (aphids and psyllids, respectively) supports the hypothesis that anti-viral immune modulation is an adaptive strategy to promote transmission and spread of diverse circulative plant pathogens. These findings open a new and unexplored avenue of research on a possible role of ISVs in insect transmission of circulative plant pathogens and/or their future use to dissect the molecular mechanisms involved in transmission.

## Acknowledgements

The authors would like to acknowledge Chad Thomas, Julie Bojanowski, Lisa Scanlon (Cornell University, Ithaca, NY) for care of the plants and aphids used in these experiments and Dr. Robert Shatters (USDA Agricultural Research Service, Fort Pierce, Florida) for graciously supplying the psyllids used for small RNA sequencing. This work was supported by USDA NIFA projects 2020-67013-31917 and 2015-70016-23028, the Emergency Citrus Disease Research and Extension Program project 2020-70029-33176 and USDA Agricultural Research Service project 8062-22410-007-000-D. This research was supported in part by an appointment to the Agricultural Research Service (ARS) Research Participation Program administered by the Oak Ridge Institute for Science and Education (ORISE) through an interagency agreement between the U.S. Department of Energy (DOE) and the U.S. Department of Agriculture (USDA). ORISE is managed by ORAU under DOE contract number DE-SC0014664. All opinions expressed in this paper are the author’s and do not necessarily reflect the policies and views of USDA, DOE, or ORAU/ORISE.

