## Supplementary material for "Viral and bacterial plant pathogens suppress antiviral defense against flaviviruses in their insect vectors": Table S1

**Table S1**. Oligonucleotides used in this study

| **Purpose** | **Oligonucleotide** | **5'-3' Oligo sequence** | **Length** | **Product size (bp)** | **PCR**  **conditions** |
| --- | --- | --- | --- | --- | --- |
| **Aphid specimen ID** | Mp_COI-F | GGTCAACAAATCATAAAGATATTGG | 25 | 709 | x1(94° for 5min); x35(94° for 40sec, 45° for 40sec, 72° for 1min); x1 (72° for 5 min) |
|  | Mp_COI-R | TAAACTTCAGGCTGACCAAAAAATCA | 26 |  |  |
|  | MpDNV-F | TGACAATGGGTATATTCATTGACCT | 25 | 207 |  |
|  | MpDNV-R | ATCGTGCGTCAAAAGAAACCCT | 22 |  |  |
| **sRNA** | 3’ Linker | rApp-CTGTAGGCACCATCAAT-Amine | 17 |  |  |
|  | Reverse-Transc. | Amine-GACGTGTGCTCTTCCGATCTATTGAT  GGTGCCTACA*G | 38 |  |  |
| **Detection** | MpFV-Hel-F | AGGAGCCGGTAAGATACAGC | 20 | 361 | x1(94° for 3min); x35(94° for 30sec, 55° for 30sec, 72° for 45sec); x1 (72° for 5 min) |
|  | MpFV-Hel-R | GGTGAAGTAGGTGTTGTCTGC | 21 |  |  |
|  | MpFV-RdRp-F | CTGTGTACGAGGCTGATCTCC | 21 | 471 |  |
|  | MpFV-RdRp-R | ACTACGTCGTCACCATCAACC | 21 |  |  |
| **Genome amplification** | Mp-flavi-F1 | TCC GAG TTG CAC ATC TAG ACC | 21 | 1390 | x1(94° for 5min); x35(94° for 30sec, 54° for 30sec, 72° for 1:40min); x1 (72° for 5 min) |
|  | Mp-flavi-R1 | GAC CAT GAG TAC GTT GAT GCC | 21 |  |  |
|  | Mp-flavi-F2 | TTC ATC AAG GCC CTT AAC AGC | 21 | 1435 |  |
|  | Mp-flavi-R2 | GTC GGC AAA CTT CCT CTA CG | 20 |  |  |
|  | Mp-flavi-F3 | TCT CCC AAC TAT GCT CAG ATG C | 22 | 1469 |  |
|  | Mp-flavi-R3 | CCA ATC GGC AAC AGA CAA AGG | 21 |  |  |
|  | Mp-flavi-F4 | TGT TAA GTC AGT CGT TCA TGT CG | 23 | 1412 |  |
|  | Mp-flavi-R4 | CTT TAC ACT CGC CGT AAC CG | 20 |  |  |
|  | Mp-flavi-F5 | TGG TGC ATA TAC TCG GAC TGG | 21 | 1505 |  |
|  | Mp-flavi-R5 | GCC ACC GTA ATA ACC TTA CCG | 21 |  |  |
|  | Mp-flavi-F6 | CTC TCT GAT GAC GCT AAA TGG C | 22 | 1533 |  |
|  | Mp-flavi-R6 | CGC ACC TTC ATT AAC GTC ACC | 21 |  |  |
|  | Mp-flavi-F7 | AGG CGA CTC ACC CGA TGT C | 19 | 1427 |  |
|  | Mp-flavi-R7 | ACA CAC TGA TAA CGT ACT TAC CG | 23 |  |  |
|  | Mp-flavi-F8 | CGT AGT AGC TGT CAA CGT CAC C | 22 | 1532 |  |
|  | Mp-flavi-R8 | CAG GGT TGT AGG CGT TGT CC | 20 |  |  |
|  | Mp-flavi-F9 | TGT TCA CCT GCC AAA GTT GC | 20 | 1557 |  |
|  | Mp-flavi-R9 | AAT GCC GTT GGT CAT GTT CC | 20 |  |  |
|  | Mp-flavi-F10 | GGA AGG CGT CGA TGT TGC | 18 | 1595 |  |
|  | Mp-flavi-R10 | AGT CAG TAC AAC CGA TAA TGG C | 22 |  |  |
|  | Mp-flavi-F11 | AGT TCA CGT CTT GGG CTA GG | 20 | 1504 |  |
|  | Mp-flavi-R11 | TTG TAG ACA AGG CCG AGA CC | 20 |  |  |
|  | Mp-flavi-F12 | TCT CGT TTA GGG ATG ATG TTG C | 22 | 1462 |  |
|  | Mp-flavi-R12 | TCC ATC TCA GGC TCG AAT CG | 20 |  |  |
|  | Mp-flavi-F13 | TGA TGC AGT TGT CCC TAG TCG | 21 | 1487 |  |
|  | Mp-flavi-R13 | CGT AGT CAG CAG CTA ACT TCG | 21 |  |  |
|  | Mp-flavi-F14 | CTG GAG AGA GAA GTT CGT TGC | 21 | 1455 |  |
|  | Mp-flavi-R14 | ACT AGA AGG CCG TAC TTG CG | 20 |  |  |
|  | Mp-flavi-F15 | ATA CCC AAG GAC GAC GAT GC | 20 | 1536 |  |
|  | Mp-flavi-R15 | TGGCGGACTTCAAAGCTTCC | 20 |  |  |
|  | Mp-flavi-F16 | CAACGTAGTTAGGAGTGACTTCAC | 24 | 1506 |  |
|  | Mp-flavi-R16 | TCCGCCCTTATTACCTGAACG | 21 |  |  |
|  | Mp-flavi-F17 | TGGGTAAGAGATGGTCAAAGGG | 22 | 1587 |  |
|  | Mp-flavi-R17 | GCTATGCTAGGGTTTGAGTGG | 21 |  |  |
| **RACE** | 744 | GACCACGCGACGTGTCGAVTTTTT TTTTTTTTTT | 34 |  |  |
|  | 743 | TTACTAATATCCCCCCCCCCCC | 22 |  |  |
|  | Mp-flavi-3end-F | ATGGTACCCTTCATCGACAGC | 21 | 236 |  |
|  | Mp-flavi-5end-R | CAGCGGGATATACGATCAACC | 21 | 397 |  |
