## Supplementary material for "Viral and bacterial plant pathogens suppress antiviral defense against flaviviruses in their insect vectors": Table S2

**Table S2**. Horizontal transmission of MpFV

|  |  |  |  | **MpFV-polymerase (RdRp)** | | | **MpFV-Helicase (Hel)** | | |
| --- | --- | --- | --- | --- | --- | --- | --- | --- | --- |
|  |  |  |  | **RT-PCR Positive** | | **Transmission** | **RT-PCR Positive** | | **Transmission** |
| **Replicate 1** |  | **Aphid species** | **Collected** | **RdRp** | **Positive** | **%** | **Hel** | **Positive** | **%** |
|  | After AAP* | *R.padi* | 10 | 1 | 1/10 | 10 | 1 | 1/10 | 10 |
|  |  | *R. maidis* | 9 | 1 | 1/9 | 11 | 1 | 1/9 | 11 |
|  |  | *A. gossypii*-cucumber | 4 | 0 | 0/4 | 0 | 0 | 0/4 | 0 |
|  |  | *A. gossypii*-cotton | 12 | 1 | 1/12 | 8 | 1 | 1/12 | 8 |
|  |  | *S. graminum* | 9 | 1 | 1/9 | 11 | 5 | 5/9 | 56 |
|  | After gut clearing | *R.padi* | 8 | 0 | 0/8 | 0 | 0 | 0/8 | 0 |
|  |  | *R. maidis* | 0 | 0 | - | - | - | - | - |
|  |  | *A. gossipy*-cucumber | 15 | 1 | 1/15 | 7 | 3 | 3/15 | 20 |
|  |  | *A. gossipy*-cotton | 9 | 1 | 1/9 | 11 | 1 | 1/9 | 11 |
|  |  | *S. graminum* | 10 | 2 | 2/10 | 20 | 2 | 2/10 | 20 |
|  |  | **Aphids tested** | **86** | **8** | **8/86** | **9** | **14** | **14/86** | **16** |
| **Replicate 2** | After AAP* | *R.padi* | 16 | 5 | 5/16 | 31.3 | 2 | 2/16 | 13 |
|  |  | *R. maidis* | 16 | 1 | 1/16 | 6.3 | 7 | 7/16 | 44 |
|  |  | *A. gossipy*-cucumber | 7 | 1 | 1/7 | 14.3 | 1 | 1/4 | 14 |
|  |  | *A. gossipy*-cotton | 7 | 1 | 1/7 | 14.3 | 0 | 1/12 | 0 |
|  |  | *S. graminum* | 7 | 2 | 2/7 | 28.6 | 0 | - | 0 |
|  | After gut clearing† | *R.padi* | 0 | 0 | - | - | - | - | - |
|  |  | *R. maidis* | 0 | 0 | - | - | - | - | - |
|  |  | *A. gossipy*-cucumber | 18 | 1 | 1/18 | 5.6 | 1 | - | 6 |
|  |  | *A. gossipy*-cotton | 14 | 3 | 3/14 | 21.4 | 0 | - | 0 |
|  |  | *S. graminum* | 7 | 1 | 1/7 | 14.3 | 0 | - | 0 |
|  |  | **Aphids tested** | **92** | **15** | **15/92** | **16.3** | **11** | **11/92** | **12** |
|  |  | **TOTAL** | **178** | **23** | **23/178** | **12.9** | **25** | **25/178** | **14** |

* Dead aphids collected before gut clearing period

† Aphids were transferred from artificial diets to their natural host for a 72-hour-gut clearing period. *Rhopalosiphum maidis*, *R. padi*, and *Schizaphis graminum* were moved to barley; *Aphis gossipii*-cucumber to cucumber, and *Aphis gossipii*-cotton to cotton.
