## Supplementary material for "Viral and bacterial plant pathogens suppress antiviral defense against flaviviruses in their insect vectors": Fig. S1

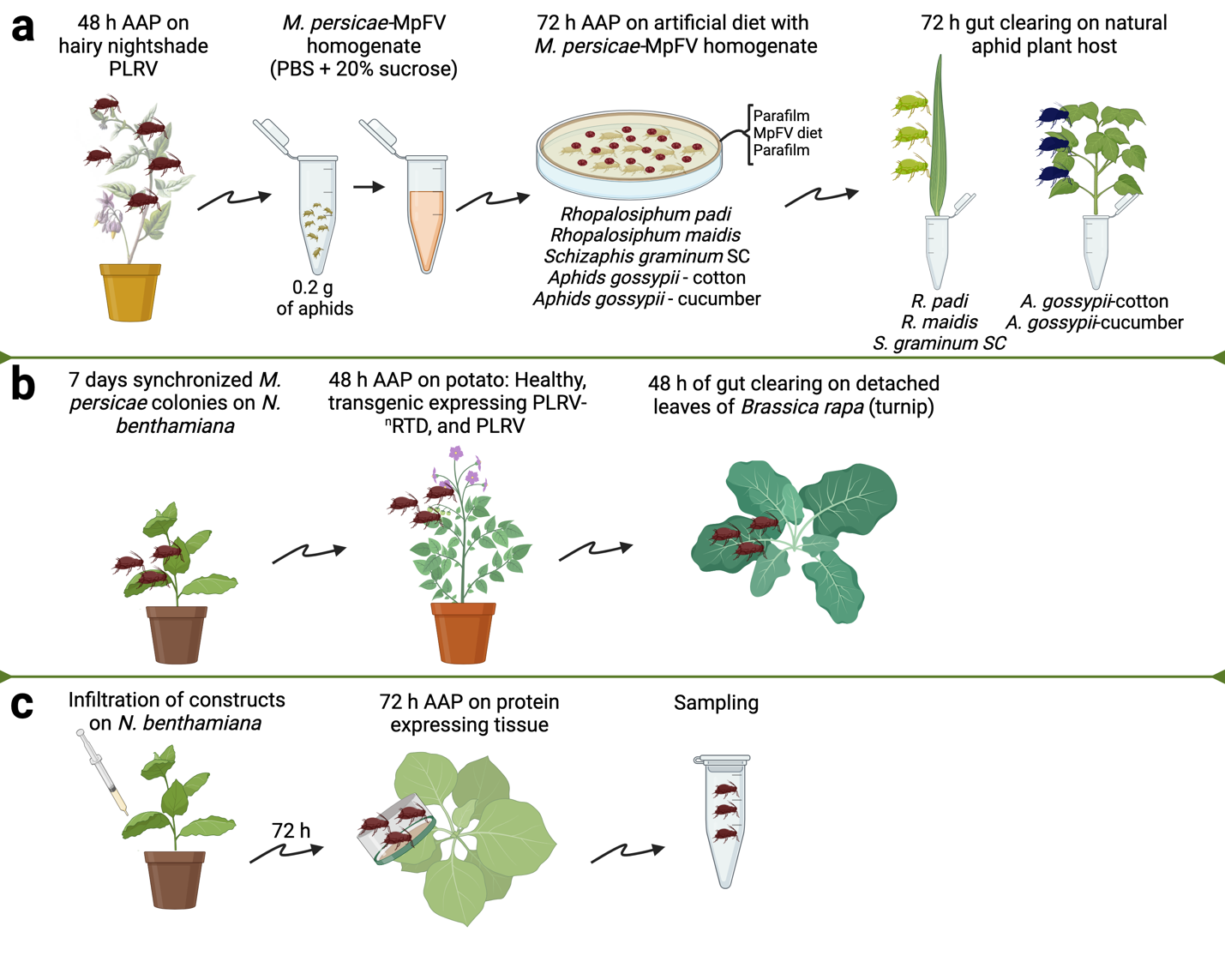

**Fig. S1.** Experimental design for aphid experiments. **(a)** Aphid species fed on MpFV artificial diet for a 72-hour acquisition access period. Then, aphids were moved to their natural host—cotton or barley—for a 72-hour gut-clearing period. To assess MpFV acquisition, RT-PCR assays were carried out on induvial aphids collected before and after the gut-clearing period. Parthenogenic aphid clone species used in this experiment include NY-*Aphis gossipii* (cotton aphid) maintained on cotton and cucumber, *Schizaphis graminium* (greenbug)*, Rhopalosiphum padi* (bind cherry-oat aphids)*,* and *R. maidis* (corn leaf aphid) maintained on barley; **(b)** Seven days synchronized Aphids fed on potato plants for 48-hour AAP followed by a gut clearing period of 48 h. Individual aphids were collected, and ddPCR was used to measure the concentration of MpFV in each treatment; **(c)** Aphid fed on transiently protein expressing *N. benthamiana* tissue for 72 hours AAP. Then, aphids were collected and tested in groups of three aphids.
