## Supplementary material for "Viral and bacterial plant pathogens suppress antiviral defense against flaviviruses in their insect vectors": Fig. S2

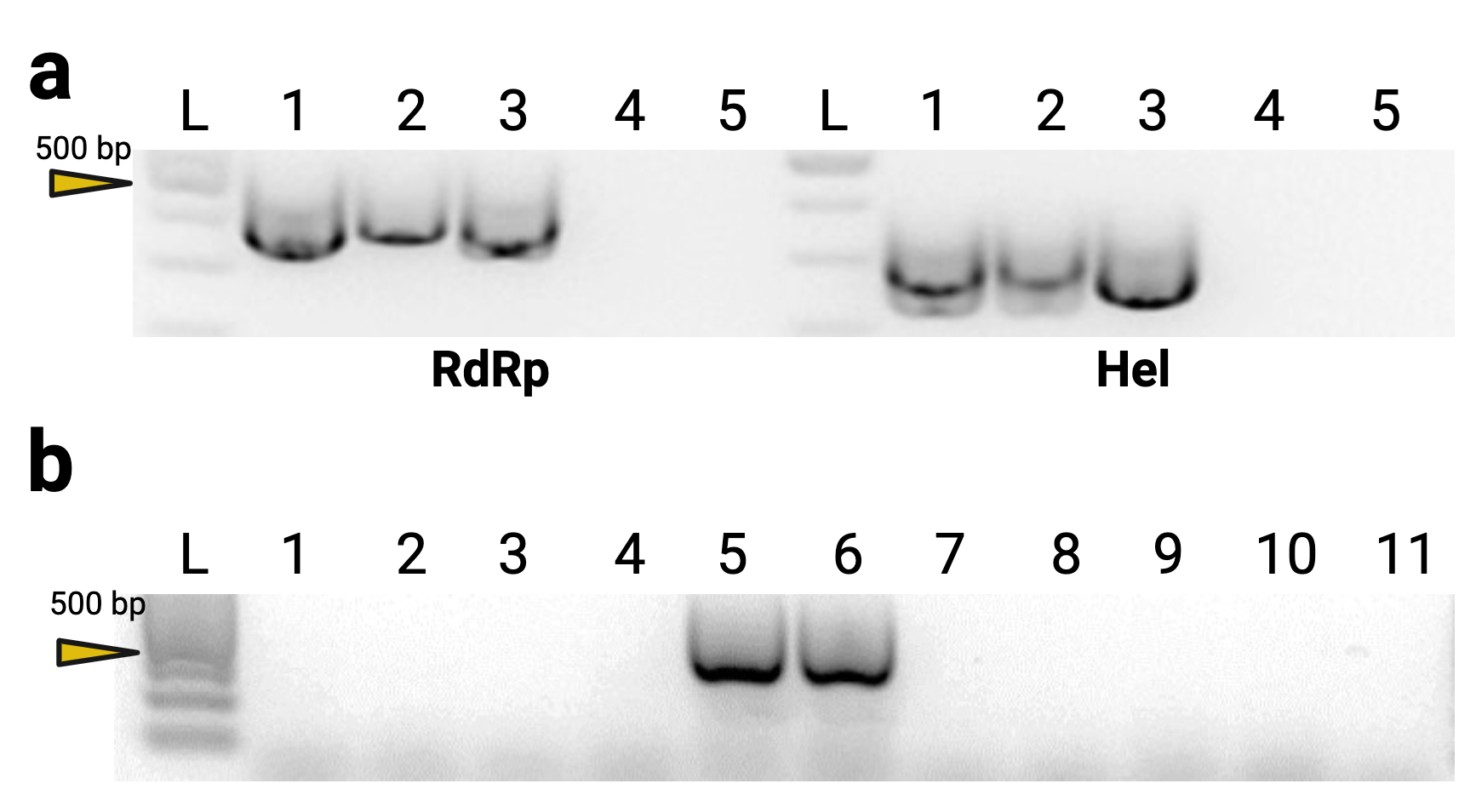

**Fig. S2.** *In vitro* validation of the presence of the novel Myzus persicae flavivirus (MpFV) infecting *Myzus persicae*. (**a**) Detection of the RNA-dependent RNA polymerase (RdRp, 471 bp) and Helicase (Hel, 361 bp) from cDNA samples prepared from RNA extracted from *M. persicae* colonies established on turnip (1), Physalis (2), and *Nicotiana benthamiana* (3). cDNA prepared from RNA from *Aphis gossypii* (4) was used a control group, and blank (5). (**b)** RT-PCR amplification of MpFV in *A. gossypii* collected from cucumber (1) and cotton from New York, Mississippi, and Alabama (2-4, respectively), *M. persicae* colonies from turnip and physalis (5 and 6), S. sp. SC (7), *Rhopalosiphum madis* (8), *R. padi* (9), and *Schizaphis graminum* (10) and blank (11).
