## Supplementary material for "Viral and bacterial plant pathogens suppress antiviral defense against flaviviruses in their insect vectors": Fig. S3

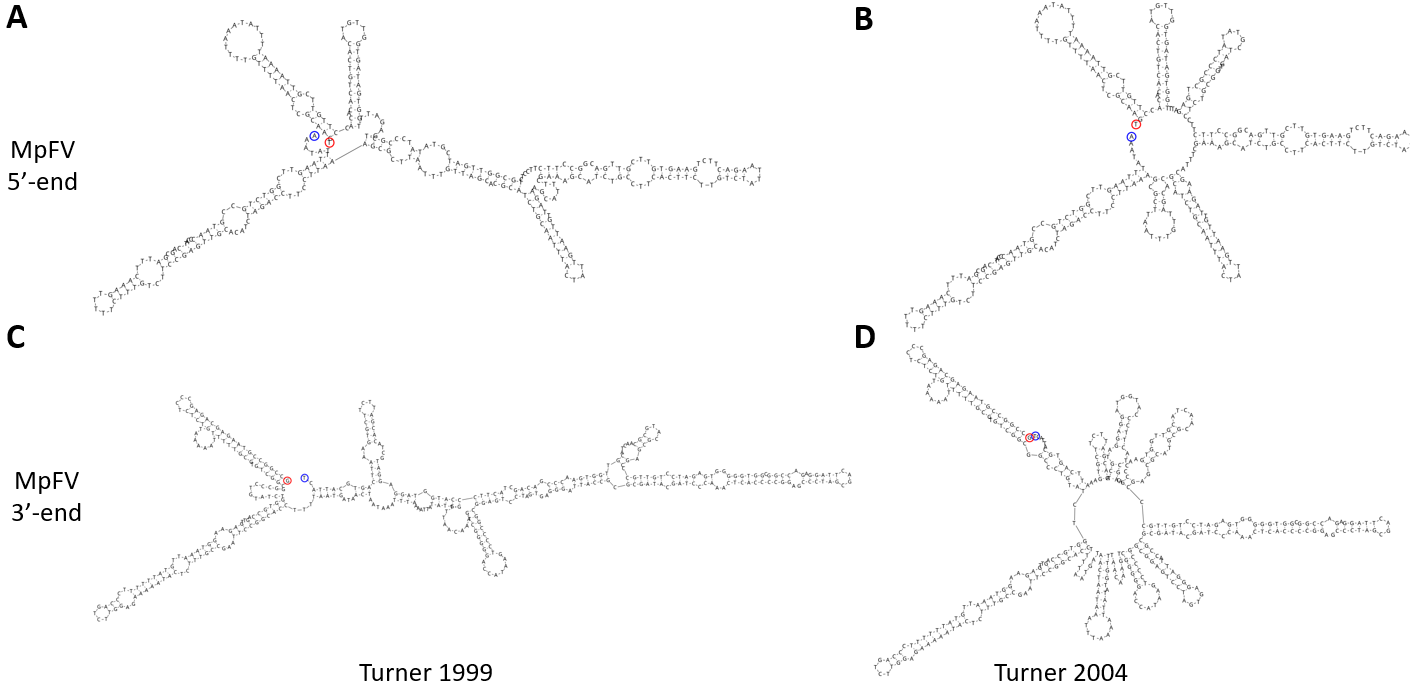

**Fig. S3**. Schematic representation of secondary structures from the untranslated regions of Myzus persicae Flavivirus (MpFV). **(a)** and **(b)** predicted MpFV 5ʹ UTR with 314 bp using Turner 1999 (left) and Turner 2004 (right) algorithms. **(c)** and **(d)** predicted MpFV 3ʹ UTR with 395 bp using Turner 1999 (left) and Turner 2004 (right) algorithms. Stem-loop structures were observed on both 5’ and 3’ ends. A temperature of 25°C was used for the predictions.
