## Supplementary material for "Viral and bacterial plant pathogens suppress antiviral defense against flaviviruses in their insect vectors": Fig. S4

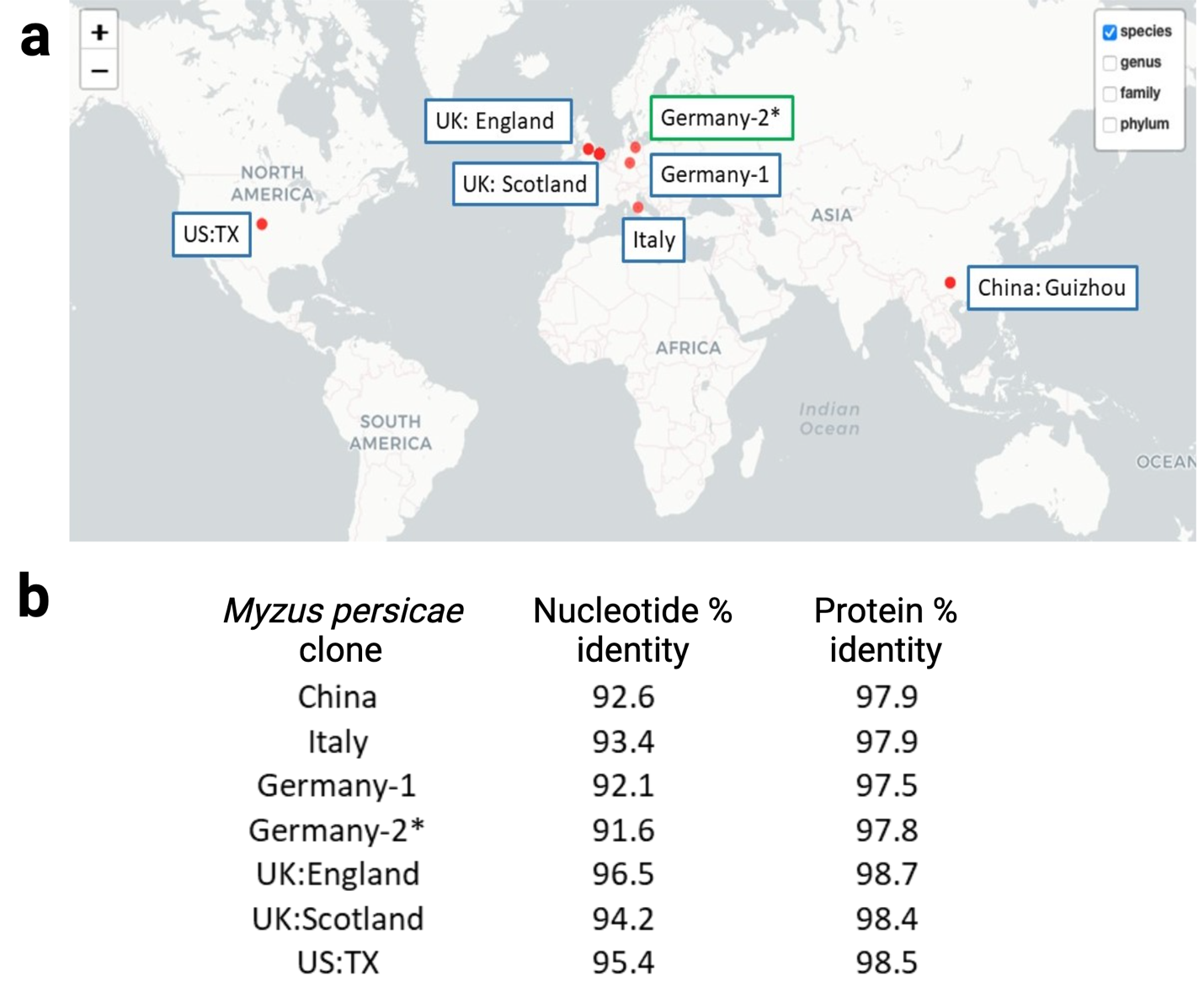

**Fig. S4.** Data mining to explore Myzus persicae Flavivirus (MpFV) diversity and tissue tropism. **(a)** Geographic distribution of MpFV on *M. persicae* samples represented blue boxes. An accession from a plant sample collected in Germany (Germany-2*), *Blitum bonus-Henricus*, is represented in green. **(b)** Nucleotide and protein identities of MpFV from data mining compared with the sequence of MpFV reported in the present study infecting MpFV.
