## Supplementary material for "Viral and bacterial plant pathogens suppress antiviral defense against flaviviruses in their insect vectors": Fig. S5

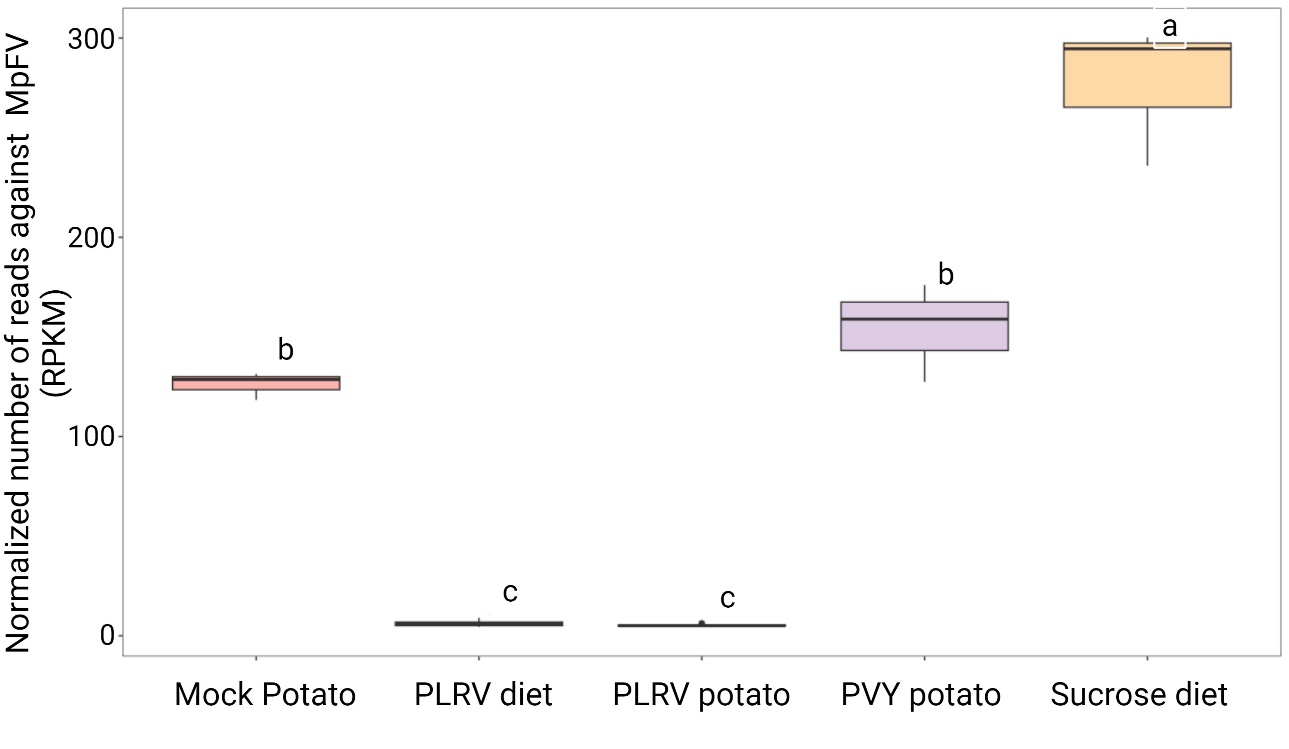

**Fig. S5**. Boxplots of the normalized number of sRNA reads per kilobase per million (RPKM) mapping against the Myzus persicae Flavivirus in aphids feeding on mock-infected potato, PLRV virions in artificial diet, systemically PLRV-infected potato, systemically PVY-infected potato and sucrose diet. All reads used for quantification matched 100% to the virus genome (no mismatches). Tukey's test determined statistically different groups (a, b, and c).
