## Supplementary material for "Viral and bacterial plant pathogens suppress antiviral defense against flaviviruses in their insect vectors": Fig. S6

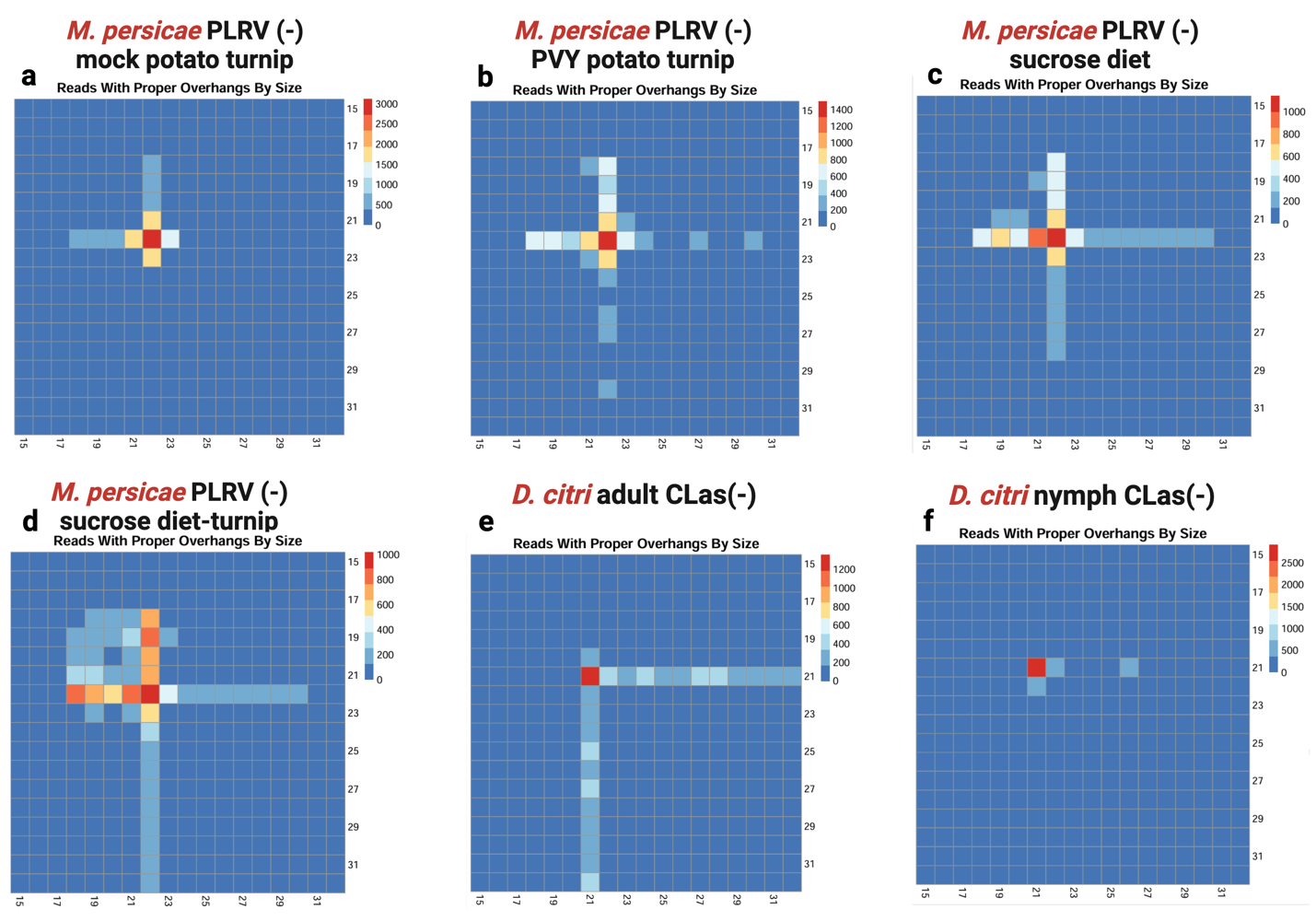

**Fig. S6**. Six heatmap matrices representing length distribution of sRNA reads from *Myzus persicae* that mapped to M. persicae Flavivirus **(a-d)** and sRNA reads from *D. citri* that mapped to DcFLV **(e & f)** which were found to overlap and have a 3’ 2nt overhang remnant of Dicer-2 cleavage. Each overlapping reads from both strands are examined and plotted as a pairwise matrix. A read length of 21 is the most abundant in *D. citri* samples (e-f) while a read length of 22 is most abundant in *M. persicae* samples (a-d). Data from only non-viruliferous or non-bacteruliferous insects were used for these analyses because there was an insufficient number of flavivirus matching-reads produced in PLRV-viruliferous aphids or *C*Las (+) psyllids.
